# The total mRNA concentration buffering system in yeast is global rather than gene-specific

**DOI:** 10.1101/2021.01.14.426689

**Authors:** José García-Martínez, Daniel A. Medina, Pablo Bellvís, Mai Sun, Patrick Cramer, Sebastián Chávez, José E. Pérez-Ortín

**Affiliations:** Instituto de Biotecnología y Biomedicina (BIOTECMED), Facultad de Biológicas, Universitat de València. C/ Dr. Moliner 50. E46100 Burjassot, Spain; Instituto de Biomedicina de Sevilla, Universidad de Sevilla-CSIC-Hospital Universitario V. del Rocío, Seville, 41012, Spain; Max Planck Institute for Biophysical Chemistry, Department of Molecular Biology, Am Fassber 11, 37077 Göttingen, Germany; Dirección de Evaluación y Acreditación, Agencia Andaluza del Conocimiento. Doña Berenguela s/n, planta 3^a^ C.P. 14006 Córdoba, Spain

**Keywords:** crosstalk, transcription, mRNA decay, yeast, aneuploidy, NMD

## Abstract

Gene expression in eukaryotes does not follow a linear process from transcription to translation and mRNA degradation. Instead it follows a circular process in which cytoplasmic mRNA decay crosstalks with nuclear transcription. In many instances this crosstalk contributes to buffer mRNA at a roughly constant concentration. Whether the mRNA buffering concept operates on the total mRNA concentration or at the gene-specific level, and if the mechanism to do so is a global or a specific one, remain unknown. Here we assessed changes in mRNA concentrations and their synthesis rates along the transcriptome of aneuploid strains of the yeast *Saccharomyces cerevisiae*. We also assessed mRNA concentrations and their synthesis rates in non sense-mediated decay (NMD) targets in euploid strains. We found that the altered synthesis rates in the genes from the aneuploid chromosome and the changes in their mRNA stabilities were not counterbalanced. In addition, the stability of NMD targets was not specifically compensated by the changes in synthesis rate. We conclude that there is no genetic compensation of NMD mRNA targets in yeast, and total mRNA buffering uses mostly a global system rather than a gene-specific one.

## Introduction

Cell homeostasis requires the total concentrations of proteins and RNAs to be remain within a certain range (Pérez-Ortín et al. 2019). In the yeast *Saccharomyces cerevisiae*, it has been shown that the total mRNA concentration ([mRNA]_t_) is buffered (Pérez-Ortín et al. 2013; Sun et al. 2013; Haimovich et al. 2013). Indeed several reports in strains with mutations in proteins related to either mRNA synthesis or decay machineries indicate the primary defect caused by mutation: a drop in global synthesis rates or degradation rates is compensated by a roughly comparable increase in the reciprocal rate (Sun et al. 2013; Haimovich et al. 2013; Timmers and Tora 2018; Begley et al. 2019). In principle, buffering can also apply to specific mRNAs or groups of functionally-related mRNAs. For instance, the existence of [mRNA]_t_ buffering in a series of yeast mutants showed that the buffering effect varies between individual mRNA species (Sun et al. 2013; Haimovich et al. 2013; Timmers and Tora 2018; Begley et al. 2019; Medina et al. 2014; García-Molinero et al. 2018). Yet despite the indirect effects of mutations, it is still difficult to unequivocally conclude whether gene-specific buffering exists or not.

In mammalian cells, the attenuation of global transcription leads to the widespread stabilization of mRNAs (Helenius et al., 2011; Slobodin et al., 2020) which, thus, demonstrates the existence of [mRNA]_t_ buffering in higher eukaryotes (Hartenian and Glaunsinger 2019). In these organisms, specific mRNA buffering has also been demonstrated in one particular case called genetic compensation, where destabilization of some defective mRNAs by the non sense-mediated decay (NMD) pathway is partially compensated by a rise in the synthesis rates of sequence-related genes (Ma et al. 2019; El-Brolosy et al. 2019).

Global or gene-specific buffering is likely driven by distinct molecular mechanisms, and the purpose of each process might be completely different. Total mRNA buffering may have evolved to keep [mRNA]_t_ within the physiological limits that maintain processes in the cytoplasm efficient, e.g., translation (Pérez-Ortín et al. 2019; Lin and Amir 2018), and to also maintain normal physico-chemical cytoplasm behavior and the solubility of cellular proteins, which very much depend on RNA concentration (Tauber et al. 2020; Aarum et al, 2020). However, the purpose of gene-specific mRNA buffering is likely an entirely different one. For instance, it could be used to transiently adjust transcriptional response timing. In fact it has been shown that the genes with or without antisense transcription have different average mRNA half-lives, and antisense transcription inactivation can lead to an increase in sense synthesis rate and a decrease in half life, which would help to make the mRNA level (or mRNA amount) constant (Brown et al. 2018). In this case, the specific buffering effect is obtained through chromatin signatures, which are established by antisense transcription and affect the initiation and elongation of sense transcription. The influence of transcription elongation on mRNA stability has been recently established (Begley et al. 2019; 2020; Fischer et al., 2020). Hence, the control by antisense transcription, which reduces production and increases stability, but maintains the same final transcript level, can be a regulatory process for specific genes and a useful one when rapidly varying conditions are expected (Brown et al. 2018).

[mRNA]_t_ buffering is based on the crosstalk between transcription and mRNA decay machineries (Sun et al. 2013; Haimovich et al. 2013; Das et al. 2017). This crosstalk functions from the nucleus to the cytoplasm (direct) (Dahan and Choder 2013), and from the cytoplasm to the nucleus (reverse) (Sun et al. 2013). As such, it has been suggested that the gene expression process in eukaryotes is circular (Haimovich et al. 2013). Direct crosstalk may occur by either mRNA imprinting (Choder 2011) with RNA-binding proteins (RBP) or the co-transcriptional methylation of specific nucleotides (Slobodin et al. 2020). The reverse crosstalk mechanism is less clear, but may be based on the titration of general RBPs in the cytoplasm with mRNA molecules. Titration by mRNAs limits the number of RBP molecules that are imported back to the nucleus and, therefore, senses [mRNA]_t_ (Gilbertson et al. 2018; Schmid and Jensen 2018; Hartenian and Glausinger 2019). However, these studies did not conclusively answer the question as to whether [mRNA]_t_ buffering is the additive consequence of gene-specific regulations of many individual mRNA species, or if it operates as a global mechanism.

Here we used the model eukaryote *S. cerevisiae* to investigate whether widespread gene-specific mRNA buffering exists. To achieve this, we employed a set of aneuploid strains with either excess of or a defect in a single copy of a chromosome. This allowed us to assess if the presumed increase/decrease in gene transcription of the genes belonging to that particular chromosome would provoke a compensatory effect on their mRNAs stabilities. We also studied NMD targets in wild-type and *upf1* mutant euploid strains to check if an increase in the mRNA stabilities of a selected group of mRNAs would provoke a specific compensatory effect on their synthesis rates. We conclude from our results that, at least in this unicellular eukaryote, most genes do not undergo gene-specific mRNA buffering, which seems mostly a global process.

## Results

For our search to find the mechanism behind [mRNA]_t_ buffering, we designed two strategies to test the existence of gene-specific mRNA buffering in the yeast *S. cerevisiae*. Both are based on the idea that if one of the two buffering parameters, synthesis rates or half lives, of a selected group of non functionally related genes changes in relation to the whole genome, it will cause a similar relative change in its mRNA levels unless a gene-specific crosstalk buffers it by acting on the reciprocal parameter.

For the synthesis rate change, we employed stable aneuploid strains in which a single chromosome had a different copy number because this would bring about a change in the synthesis rates only in the set of genes contained in that chromosome. To illustrate half life alteration, we analyzed the mRNAs affected by the NMD pathway that destabilized mRNAs with premature stop codons and other specific sequence features (Celik et al. 2017).

### Transcriptomic study of a set of aneuploid yeast strains

In order to investigate the alteration of synthesis rates in a non functionally related group of genes, we took advantage of the fact that some yeast strains have an extra copy of one chromosome (disomic/trisomic) in haploid/diploid backgrounds, or only one copy of a chromosome (monosomic) in diploid strains. In these strains, we expect an increase or decrease in the synthesis rate, respectively, of the genes belonging to the aneuploid chromosome in relation to the rest of the genome. The buffering effect cannot be investigated by simply looking at the mRNA level, as other researchers have previously done (Torres et al 2007; Hose et al. 2015), because of the possible influence of both synthesis rate and half life on the actual mRNA level. The growth phenotype of many single-gene mutant yeast strains is partially suppressed by the over-/underexpression of a set of genes contained in a given chromosome (Hughes et al. 2000). If the average synthesis rate and mRNA amount in the aneuploid chromosome vary in parallel in relation to the other chromosomes (e.g. no change in average half life), then we can reject the gene-specific buffering hypothesis.

We first employed a series of mutant strains that had been determined as aneuploids by two different genomic methods in two separate laboratories: Genomic Run-on (GRO, Haimovich et al. 2013) and a comparative Dynamic Transcriptome Analysis (cDTA, Sun et al. 2013). All these mutant strains were initially assumed to be euploids. After discovering that some of them had an additional copy of one chromosome or were diploids with a single copy of one chromosome it was necessary, in the original study, to remake them and check that they were euploids (Sun et al. 2013). In the present study, we used the initial aneuploid strains to analyze their mRNA amounts, synthesis rates and half lives. As seen in Figure 1A, all the haploid strains with a disomy displayed an average increase in the synthesis rate of about 1.71-fold (except for mutant *edc1*) in relation to the whole genome. This was similar to that of the mRNA amount (1.76-fold) for aneuploid chromosomes as regards the whole transcriptome (Table 1). The diploid strains with one monosomy showed an average 0.56-fold decrease in both the synthesis rate and mRNA amount for aneuploid chromosomes (Figure 1B-C and Table 1). Thus we concluded that no change occurred in the half lives of the mRNAs of the chromosome with an altered ploidy.

**Table 1.** List and averages of the ratios of the medians for synthesis rates (SR), mRNA amounts (RA) and mRNA half-lives (HL) of the genes from the copy number altered chromosome vs the rest of the genome in a series of aneuploid strains. Ploidy of each mutant (n or 2n) is indicated.

|  | Mutant n | Duplicated chromosome | SR change | mRNA change | HL change |
| --- | --- | --- | --- | --- | --- |
| *Sun et al. 2013* | not4 | II | 1.729 | 1.691 | 0.978 |
|  | rrp47 | IX | 1.701 | 1.845 | 1.085 |
|  | edc1 | XI | 1.321 | 1.382 | 1.046 |
|  | pat1 | II | 1.778 | 1.860 | 1.046 |
|  | rrp6 | XII | 1.553 | 1.670 | 1.075 |
|  | pop2 | VIII | 1.750 | 1.819 | 1.040 |
|  | trf5 | IX | 1.741 | 1.770 | 1.054 |
|  | xrn1 | XI | 1.741 | 1.865 | 1.071 |
|  | Mutant 2n | Not duplicated chromosome | SR change | mRNA change | HL change |
| *Sun et al. 2013* | cbp20 | III | 0.535 | 0.540 | 1.010 |
|  | hpr1 | IX | 0.531 | 0.514 | 0.969 |
| *Medina et al. 2014* | xrn1 | III | 0.583 | 0.602 | 1.033 |
|  | Aneuploid n strain | Duplicated chromosome | SR change | mRNA change | HL change |
| This work. A. Amon's collection | A6863 | I | 1.710 | 1.498 | 0.876 |
|  | A13628 | VIII | 1.578 | 1.787 | 1.132 |
|  | Aneuploid 2n strain | Duplicated chromosome | SR change | mRNA change | HL change |
|  | A18345 | I | 1.471 | 1.168 | 0.794 |
|  | A18349 | XIV | 1.517 | 1.466 | 0.966 |
|  | A18346 | V | 1.254 | 1.406 | 1.121 |
|  | A18347 | VIII | 1.302 | 1.296 | 0.996 |
| **AVERAGES** |  |  | **SR change** | **mRNA change** | **HL change** |
| n+1 strains (mutants) |  |  | 1.71±0.07 | 1.76±0.08 | 1.05±0.03 |
| 2n-1 strains (mutants) |  |  | 0.53±0.003 | 0.53±0.02 | 1.01±0.03 |
| n+1 strains (A. Amon) |  |  | 1.64±0.09 | 1.64±0.20 | 1.00±0.18 |
| 2n+1 strains (A. Amon) |  |  | 1.39±0.13 | 1.33±0.13 | 0.97±0.13 |

**Figure 1.**
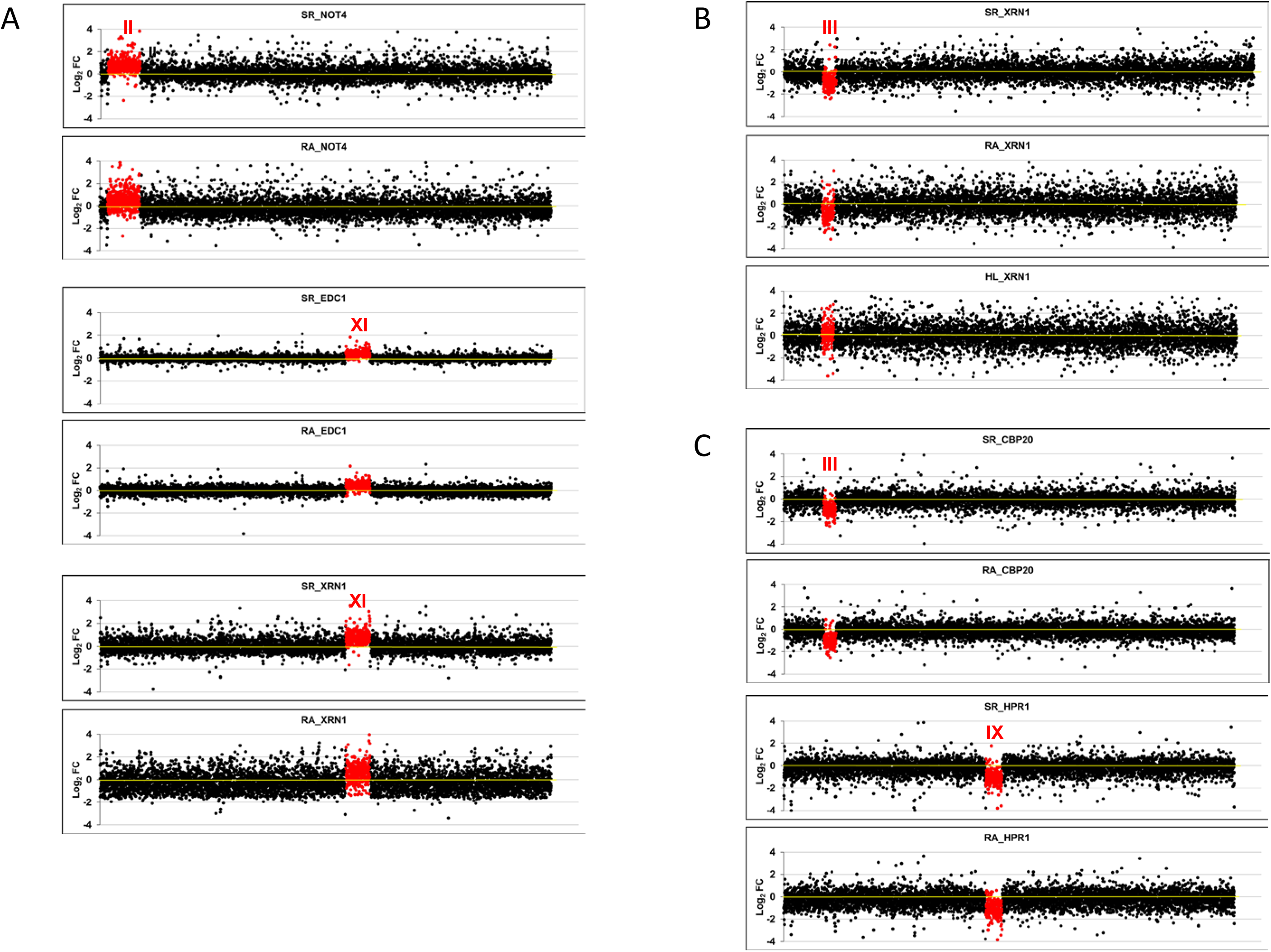
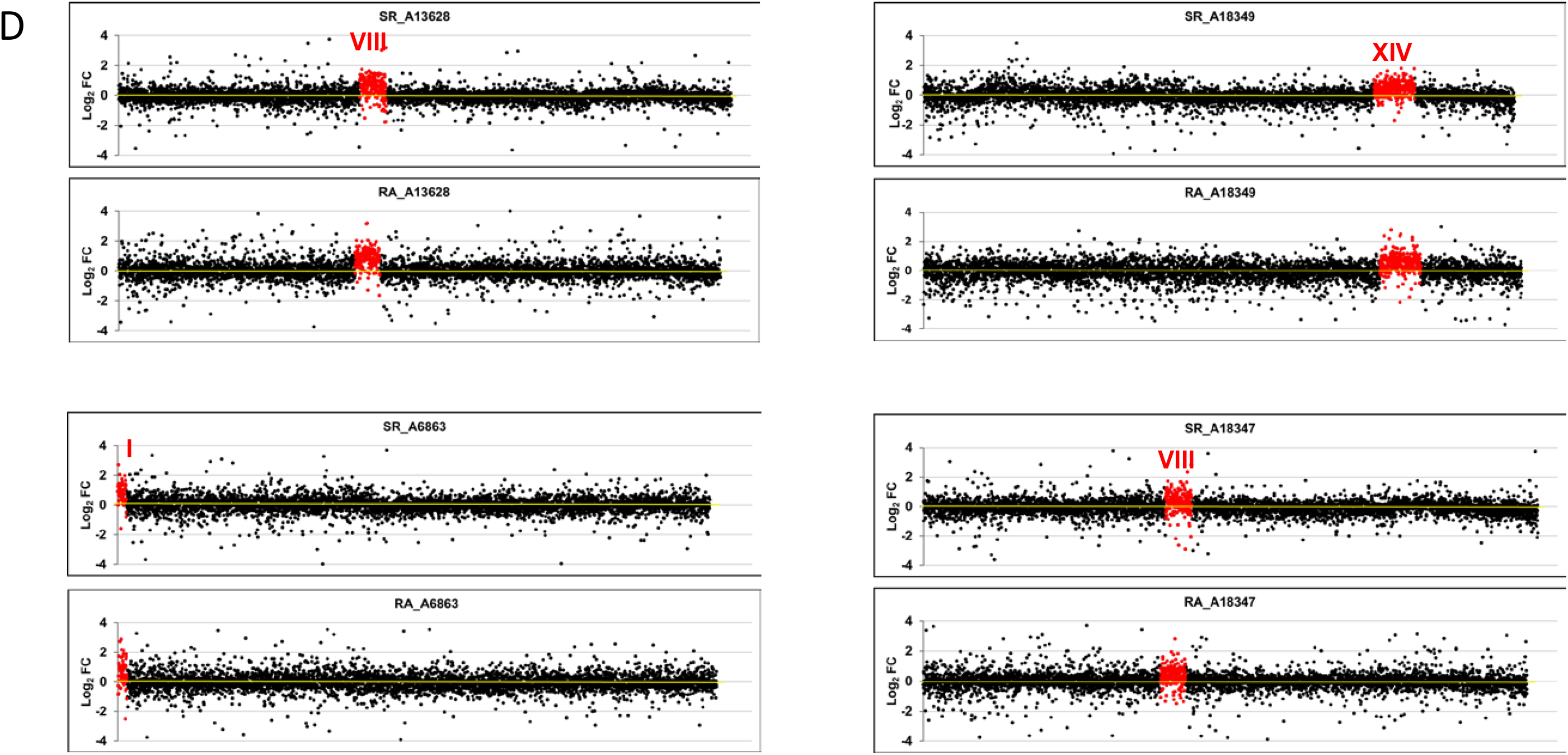
Transcriptomic analysis of aneuploid strains. We performed GRO or a cDTA analysis of aneuploid strains compared to their wild-type strain (BY4741). Then we compared the fold changes of the synthesis rates (SR), mRNA amounts (RA) and mRNA half-lives (HL) of the genes aligned from the left to right telomeres in each chromosome ordered from chromosome I (left) to chromosome XVI (right) on a log_2_ scale. The genes from the aneuploid chromosome (marked as a roman numeral) are highlighted in red. A) The results of the cDTA analysis of haploid strains *not4, edc1* and *xrn1*. B) The results of the GRO analysis of the *xrn1* diploid strain from (Haimovich et al., 2013) showing the SR, RA and HL data. The calculated HLs from the RA and SR data indicate how chromosome III has no average HL that differs from the other chromosomes. C) The results of the cDTA analysis of diploid strains *cbp20* and *hpr1*. D) The results of the GRO analysis of a set of diploid (left) and haploid (right) strains from A. Amon’s aneuploid collection (Torres et al. 2007). Note that the HL plot is shown only in the *xrn1* mutant for simplification. In all cases, the absence of chromosome-specific variation in HL (see Table 1) indicates the absence of a gene- specific buffering system. Data used in these figures are available in supplementary tables S2 and S3.

The previous strains were all mutants in the genes associated with mRNA decay. Despite the fact that the deleted gene had no direct relation to most of the genes in the altered copy chromosome, an indirect effect of the mutation occurring on the buffering phenomenon could be argued. To address this issue, we carried out another experiment with a series of stable aneuploids with no known mutations requiring compensation. We specifically utilized a series of haploid or diploid strains with an extra chromosome constructed in A. Amon’s laboratory (Hose et al. 2015). Once again, and as seen in Figure 1D, the higher synthesis rate values (1.64 for the disomic chromosome in haploid and 1.39-fold for the trisomic chromosome in diploid strains) were comparable to those for the increase in mRNA amount: 1.64- and 1.33-fold, respectively (Table 1). This indicated that no specific buffering for mRNAs with an increased synthesis rate existed.

From these experiments, we concluded that when the copy number of a group of genes was altered compared to the rest of the genome, the synthesis rates increased or decreased accordingly. This increase or decrease took place to a lesser extent than we expected (see the Discussion). As this change in the average synthesis rate was comparable to that observed in the average mRNA amount in all cases, we concluded that no specific mechanism existed in yeast to detect altered levels of individual mRNAs and to correct them by half life change compensation.

### Alteration of mRNA stability via NMD: effect of a premature STOP codon

We first investigated if the NMD pathway is able to direct the specific crosstalk that buffers the increased decay of the mRNAs containing a premature stop codon (PTC) in the same way as in metazoa (Ma et al. 2019; El-Brolosy et al. 2019). To check whether this behavior was also present in yeast, we engineered a copy of a fusion gene with or without a PTC (Fig. 2A). We measured the stability of both mRNAs and found a significant decrease in the half life of the allele containing the PTC (Fig. 2B, left panel). As an internal control, we measured the stability of the *GAL1* gene, which is driven by the same promoter, and found no significant change (Fig 2B, left panel). To further confirm that the diminished stability of the PTC-containing mRNA was due to the action of the NMD system, we repeated the experiments with an *upf1*Δ mutant, which lacked one of the fundamental NMD machinery factors. As expected, the mRNA stability of the PTC-containing allele significantly increased, and no changes were detected in the internal *GAL1* control (Fig 2B, right panel).

**Figure 2.**
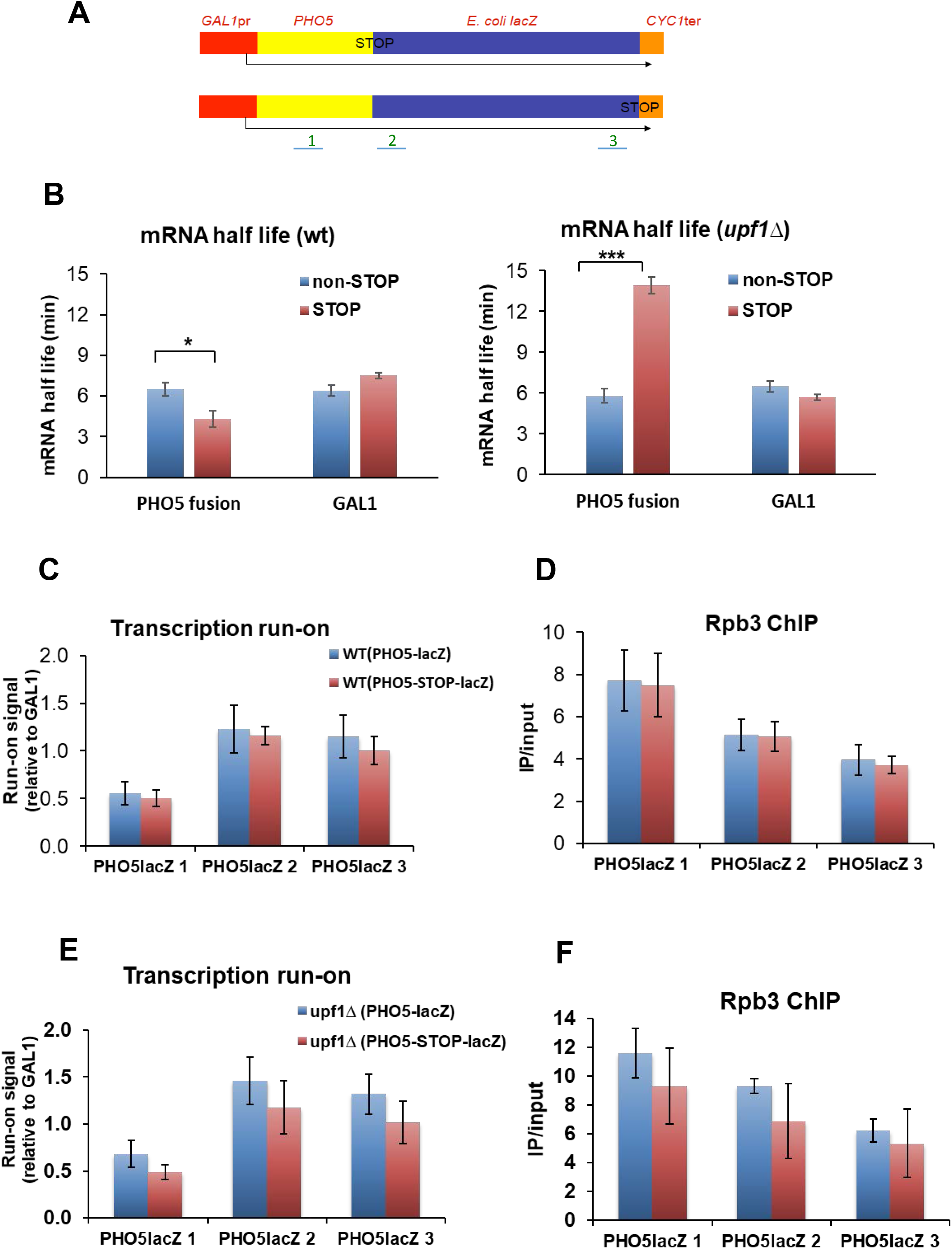
Effect of the premature stop codon (PTC) on mRNA stability and the synthesis rate. A). Scheme of the two plasmidic gene fusions used for these experiments. The *GAL1* promoter directs the synthesis of an mRNA that contains the yeast *PHO5* and *E. coli lacZ* open reading frames fused in-frame until the natural *lacZ* stop (bottom), or containing a PTC between the two ORFs (top). The *CYC1* terminator is placed at the end of constructs. The three probes (1-3) used for transcription run-on (TRO) and RNA polymerase II chromatin immunoprecipitation (ChIP) with the anti- Rbp3 antibody are shown. B) The mRNA half-life (HL) was determined by the transcription shut-off of the *GAL1* promoter by changing cells from galactose to glucose-containing media, and a northern blot of the extracted RNAs, which was successively tested with probes *PHO5, GAL1* and 18S rRNA. Note that the presence of a PTC (pink bars) provokes a one third reduction in the HL in the fusion transcript, but no decrease in the endogenous *GAL1* mRNA in the wild-type cells. Conversely, the PTC-containing fusion transcript was significantly stabilized in an *upf1* mutant. The greater stability of the PTC-containing transcript *versus* the non containing one in *upf1*Δ could be related to the much shorter translated region of the former. The shown HL values correspond to the average of at least three independent experiments. C) The TRO experiment with the wild-type strain shows no significant difference in elongating RNA pol II (SR) in the fusion genes containing (pink bars) a PTC, or not (blue bars). Values were presented after normalizing to the plasmid copy number measured by Q-PCR. The results were similar using probes 1-3. D) ChIP using anti-Rpb3 shows similar results as the TRO experiment. E) The TRO experiment in the *upf1* mutant shows that there is no significant difference in elongating RNA pol II (SR) on fusion genes containing (pink bars), or not (blue bars), a PTC. In fact the SR is slightly lower in the PTC-containing mRNA, but not statistically significant. The results were similar using probes 1-3. F) ChIP employing anti-Rpb3 in the *upf1* strain gave similar results to the TRO experiment. Bars represent the average and SD of three independent replicates of the experiment. The statistical significance of the differences between the averages of the indicated samples was estimated using a two-tailed Sudent’s t-test (* means *p* < 0.01; *** means *p* < 0.0001).

Next we assessed whether the RNA pol II synthesis rate changed in the PTC-containing allele in parallel to mRNA stability. We measured the synthesis rate by transcription run-on (TRO, measuring active elongating RNA pol II; Fig. 2C) and chromatin immunoprecipitation (ChIP, measuring total RNA pol II; Fig. 2D). The results showed that the synthesis rate in the two alleles was similar, which indicates the absence of a specific compensation for a PTC-containing mRNA in yeast. No significant differences in the synthesis rate between the two alleles were found in the *upf1*Δ background either (Fig. 2E-F). These results revealed that a change in the stability of a specific mRNA did not bring about the increase in its coding gene synthesis rate.

### Crosstalk for post-transcriptional regulons: study of the *upf1*Δ mutant

We then used the NMD system to determine if specific buffering regulated non engineered mRNAs at the decay level. We used the same *upf1*Δ mutant as that employed in the previous experiment. Upf1 is known to be cytoplasmatic in yeast, where it degrades both mRNAs with PTCs and normal-appearance mRNAs. These non-PTC containing mRNAs have either a tendency to increase out-of-frame translation or have a lower average codon optimality, plus a biased distribution pattern of non optimal codons. This set contains about 900 targets, shared with factors Upf2/Upf3 (Celik et al., 2017; He and Jacobson 2015). This group of mRNAs comprises genes with no evident functional relation. By GRO, we first analyzed the global yeast transcriptome behavior in the *upf1*Δ mutant by comparing the relative changes in both the half life and synthesis rate (Figure 3A). As expected, we found a group of mRNAs (mostly coinciding with known Upf1 targets; not shown), in which half lives were much longer. However, the change in the synthesis rate of these genes was minor, and led to a significant and parallel increase in mRNA amount (Figure 3B, left bars). To verify these results, we next studied the behavior of the whole set of previously known Upf1 targets (Celik et al. 2017) by analyzing these genes in comparison with the rest of the *upf1*Δ mutant transcriptome. Figure 3B (right bars) depicts a clear and statistically significant increase in the half lives in the targets in relation to the other mRNAs which, by not being compensated by a change in the synthesis rate, provoked a statistically significant relative increase in the mRNA amount of NMD targets. This result indicates again the absence of gene-specific buffering when the half lives from a group of genes are altered.

**Figure 3.**
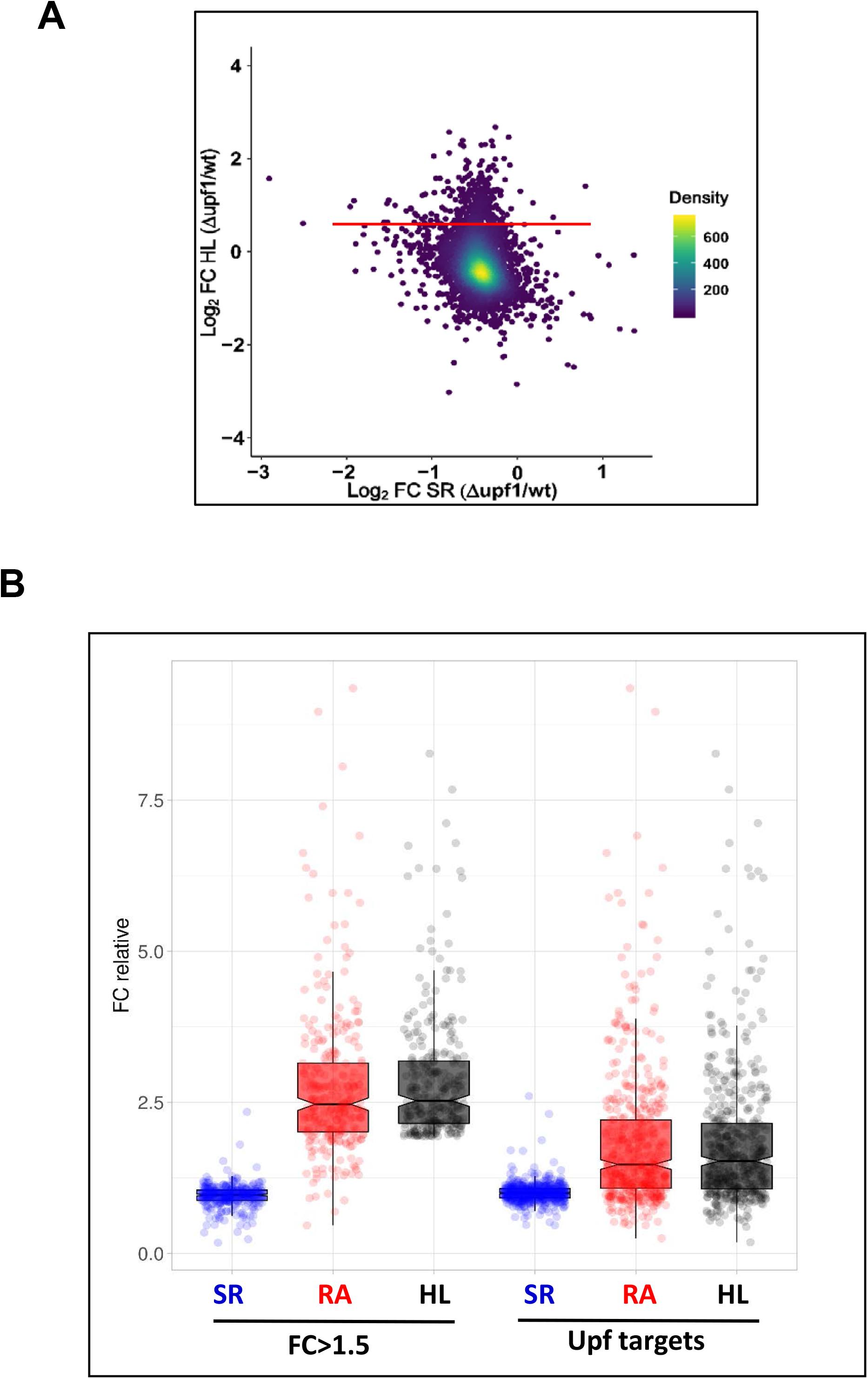
Transcriptomic analysis of the *upf1* mutant. We performed a GRO analysis of the *upf1* mutant compared to its wild-type (WT) strain (BY4741). A) Fold change (FC) in the mutant as regards the WT of half-lives (HL) vs. the FC of the synthesis rates (SR) on the log_2_ scale. The red line marks the mRNAs with FC = 1.5 in the absence of the Upf1 protein. B) Box-plot showing the fold-changes (FC) between the *upf1* mutant and the wild-type strain for the groups of mRNAs with FC>1.5 (left) and 496 Upf targets (Celik et al, 2017), right. Data are relative to the population median taken as 1. The median values for the synthesis rates (SR), mRNA amounts (RA) and mRNA half lives (HL) are shown in the plots. The experiment was run in triplicate. Data used for this figure are available in supplementary Table S4.

## Discussion

The mRNA buffering concept is often applied to the total set of mRNAs, i.e. to [mRNA]_t_, but whether it acts through a global system or the mechanism is gene-specific was previously unknown. The results obtained to date by transcriptomic methods reveal that buffering is global, albeit with major differences between individual mRNAs or between groups of mRNAs in yeast (Sun et al. 2013; Haimovich et al. 2013; Timmers and Tora 2018). This means that both global and specific mechanisms may exist at the same time.

In order to determine the possible existence of global or gene-specific processes for mRNA buffering in yeast, we devised three experiments. The three experimental setups were based on the idea that if a specific buffering mechanism exists, it must be detectable in single genes or in a group of non functionally-related mRNAs by a compensatory specific change in either the half lives or the synthesis rates.

In our first experiment, we investigated groups of the genes belonging to a given chromosome. In the yeast genome, genes mostly occupy random positions with no functional clustering (Dujon 1996). Thus choosing strains with aneuploidies allows checks to be made of what happens with the mRNA stability of functionally unrelated transcripts when changing the chromosome copy number as regards genome content. We obtained our results with different sets of aneuploids (in mutant or wild-type strains), which showed that the average synthesis rate and mRNA level underwent parallel variations in the genes of the altered chromosome in relation to the other chromosomes. This indicated lack of gene-specific buffering.

We expected an increase of 2-fold or a decrease of 0.5-fold in the synthesis rate of the genes in aneuploid strains. However, actual average increase and decrease were lower at 1.71- and 0.56-, respectively (except for *edc1*Δ: see Table 1 and Fig. 1A-C), which has also been observed by other labs (Torres et al. 2007; Hose et al. 2015). Indeed in the set of stable aneuploids from A. Amon’s laboratory, the expected increase of the synthesis rate in haploid strains was 2-fold and 1.5-fold in diploid strains, but the actual results were lower than expected, with 1.64- and 1.39-, respectively (see Table 1 and Fig. 1D). This means that in all cases of extra chromosome copies, the obtained increases in the synthesis rate were between 22% and 35% lower than expected. If the copy number of a chromosome lowered in a diploid strain, the observed decrease was also 12% lower than expected. We hypothesize this because the titration of specific transcription factors becomes limiting when their targets increase in copy number. Alternatively, these yeast cultures could have a leaking percentage of euploid cells (Torres et al. 2016), which would make the average lower than expected. In any case, the differences of the observed changes in mRNA levels (1.76- and 0.56-, see Table 1) *versus* the expected ones (2-fold and 0.5-fold by assuming a trivial translation of the increase in the gene copy number to the final mRNA level) reinforces the need of evaluating both the synthesis rates and half lives of all the mRNAs to verify the existence or nonexistence of gene-specific buffering.

In our second experiment, we compared two engineered versions of a transcription unit, one of which was an NMD target via the inclusion of a PTC. The stability of the two resulting mRNAs clearly differed, but they had identical synthesis rates.

In our third experiment, we assessed how a group of non functionally-related mRNAs, controlled by NMD (Celik et al. 2017), performed when the main NMD factor, Upf1, was lacking. Once again, our results did not support the existence of gene-specific reverse crosstalk based on NMD. Interestingly, this contrasts with what has been observed in metazoa, where NMD mediates a mechanism (NITC) of transcriptional compensation for specifically destabilized mRNAs (El-Brolosy et al 2019). This difference could be due to the much shorter genes and fewer introns in yeast compared to metazoa. Because of this, the probability of transcription and pre-mRNA splicing errors that cause non functional mRNAs with PTCs is much lower in yeast and, consequently, there may be no need for compensation at this level.

Our results supported a model in which [mRNA]_t_ buffering did not result from adding individual gene-specific buffering phenomena but, instead, was the product of the global crosstalk between mRNA decay and gene transcription machineries using direct and reverse branches (Figure 4). Based on these results, we propose a model for transcription-degradation crosstalk grounded on the co-transcriptional imprinting of mRNAs with general factors, such as RNA pol II subunits (Rpb4/7), mRNA decay factors (Xrn1, Ccr4-NOT), or others. This model is similar to those proposed by Gilbertson et al. (2018) and Schmid and Jennsen (2018) for the nuclear decay of pre-mRNAs and polyA-binding proteins (PABP, Nab2). In the event of [mRNA]_t_ buffering, the mRNA decay-related proteins would act as buffering factors and would bind pre-mRNA during transcription elongation, and not necessarily in the poly(A) tail. They would travel from the nucleus to the cytoplasm only when bound to mRNA, and would then return to the nucleus by active nuclear import after being released from their targets. Buffering factors can be part of liquid droplets (Tauber et al, 2020) formed by mRNAs where they can also be in association/dissociation equilibrium. This model would explain the [mRNA]_t_ buffering in situations in which the synthesis rate or DR would globally increase or decrease compensatorily to one another.

**Figure 4.**
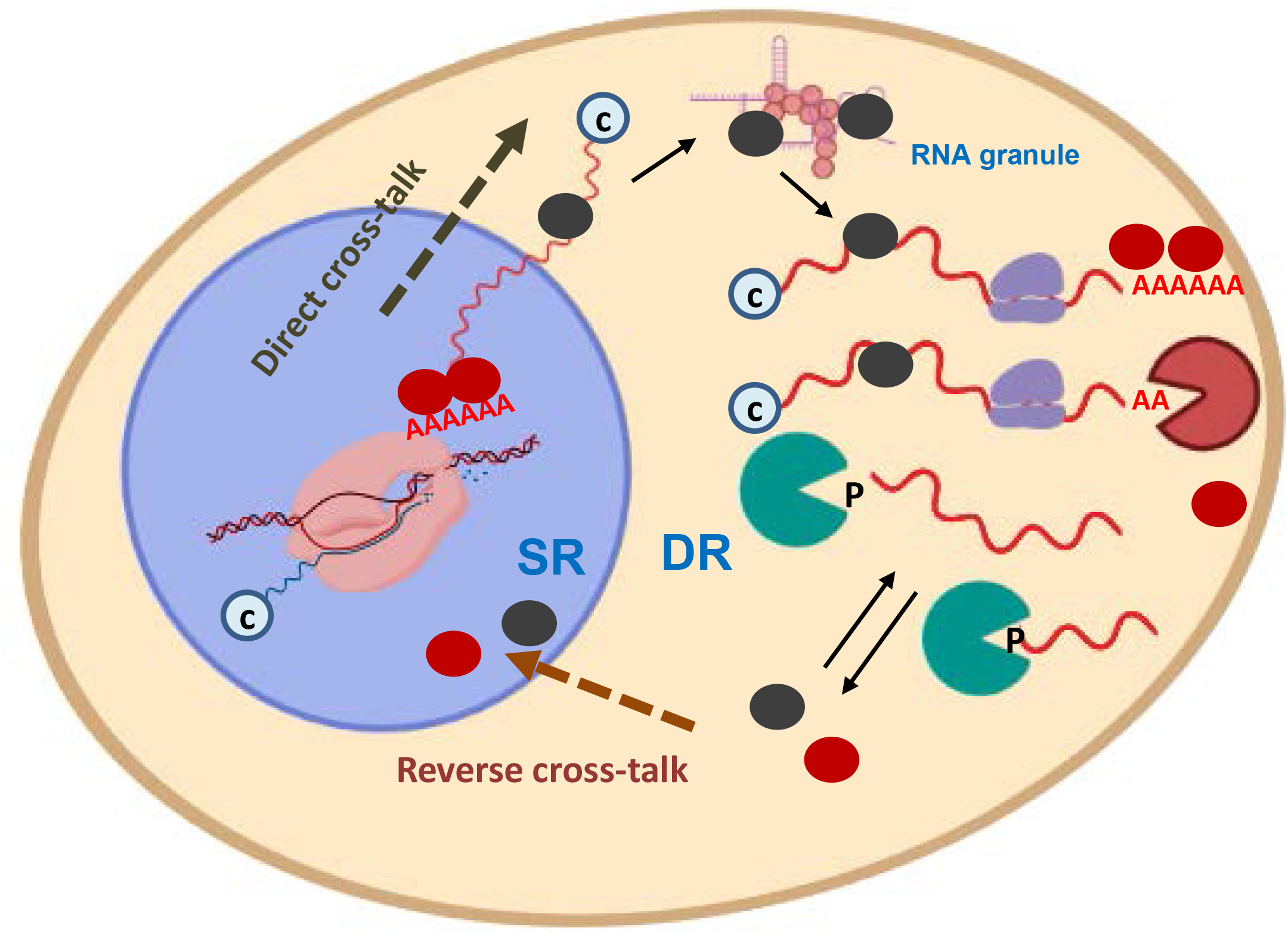
Model for mRNA buffering mediated by the nucleo-cytoplasmic shuttling of RNA-binding factors. Cartoon model showing the crosstalk pathways from the nucleus to the cytoplasm (direct), and from the cytoplasm to the nucleus (reverse). Buffering factors (BF) (colored ovals) shuttle between the cytoplasm and the nucleus. They are imported to the nucleus when they become free of mRNAs, either when mRNAs are degraded (bottom) by decay machineries (green and orange *pacman*) or before an equilibrium is struck between the bound and free states (top). BF can travel back to the cytoplasm by co-transcriptional imprinting in either mRNA sequences (black ovals) or poly(A) tails (red ovals). BFs can accompany mRNAs to p-bodies or other kinds of liquid droplets were they become separated from the soluble and translatable pool by a phase transition. Synthesis (SR) and degradation (DR) rates are adjusted for [mRNA]_t_ buffering due to the stimulating activity of the nuclear BF in RNA pol II transcription and the (direct or indirect) relation of BF to mRNA degradation in the cytoplasm. mRNAs may have a cap (c) or phosphate (P) on their 5’- end, and may be translated by the ribosomes in the cytoplasm. Figure created with the help of *BioRender*.

Transcription-mRNA decay crosstalk has been observed in mutants lacking RBPs, which may act as buffering factors (Sun et al. 2013), including Rpb4/7 (Das et al. 2017; Dahan and Choder 2013; Choder 2011), decaysome (Haimovich et al. 2013), Ccr4-NOT (Das et al. 2017; Rambout et al. 2016; Reese 2013; Collart and Panasenko 2017) and polyA-binding proteins (Schmid and Jensen 2018; Hartenian and Glausinger 2019). This suggests that mRNA buffering is a very robust process that involves many redundant pathways. Accordingly, we recently found that Xrn1 and Ccr4, two decay factors also capable of influencing transcription elongation, act in parallel (Begley et al. 2019).

To summarize, our results support the existence of global systems involved in the compensation of total mRNA levels to maintain [mRNA]_t_ within the required ribostasis limits. This is likely because the precise balance of molecules and macromolecules required for the correct functioning of cytoplasmic cell processes, including translation, albeit perhaps not exclusively (Pérez-Ortín et al. 2019). The existence of a global buffering system does not impede the particular regulation of specific mRNAs because the impact of variation in a few mRNA species would be very low in [mRNA]_t_. Our results obtained with aneuploid strains and the NMD system excluded the possibility of an all-purpose general mechanism based on specific mRNA recognition, and one devoted to compensate alterations in the proportion of cytoplasmic mRNAs species one by one.

## Materials & Methods

### Yeast strains and culture media

Yeast cells were grown in YPD (1% yeast extract, 2% peptone, 2% glucose) or YPGal (1% yeast extract, 2% peptone, 2% galactose) media at 30°C. Pre-cultures were grown overnight in 250 mL flasks and agitated at 190 rpm. The next day, pre-cultures were diluted to OD_600_ = 0.05 and grown until an OD_600_ of ∼0.5 was reached. Cells were recovered by centrifugation and flash-frozen in liquid nitrogen. Yeast strains are listed in Supplementary Table S1. For the experiments described in Figure 2, BY4741 (wt) and Y06214 (*upf1*) strains were transformed with either centromeric plasmid pSCh212 (Morillo-Huesca et al. 2006) containing the in-frame translational *PHO5-lacZ* fusion or a derivative plasmid including a *PHO5-STOP-lacZ* fusion.

### Genomic methods

Genomic run-on (GRO) was performed as described in (García-Martinez et al. 2004) as modified in (García-Molinero et al., 2018). Briefly, GRO detects by macroarray hybridization, genome-wide, active elongating RNA pol II, whose density per gene is taken as a measurement of its synthesis rate (SR). At the same time, the protocol allows the mRNA amounts (RA) for all the genes to be measured. mRNA half-lives are calculated as RA/SR by assuming steady-state conditions for the transcriptome. GRO datasets are available at the Gene Expression Omnibus (GEO) with accession numbers: GSE29519, GSE57467 and GSE155372. cDTA was performed as described in (Sun et al. 2012; 2013). Briefly, cDTA is based on the *in vivo* incorporation of 4-thiouracil into new mRNAs for a short period. The amount of newly synthesized and total mRNAs is quantified by microarray hybridization. Thus by assuming steady-state conditions, SRs and Degradation Rates (inverse to half lives) of all the genes are measured genome-wide. The cDTA data are deposited in ArrayExpress with accession number E-MTAB-1525.

### Specific mRNA half-life calculations

In order to determine single-species mRNA half lives, we ran a transcription shut-off assay by collecting samples at 0, 5, 10 and 15 min after glucose addition. This method was the same as that described in (Begley et al. 2019), except that Northern blot hybridization was used instead of RT-PCR. half lives were estimated by calculating the time it takes for half the initial amount of mRNA to be degraded. mRNA extraction and Northern blots were carried out following the protocols detailed in (Morillo-Huesca et al. 2006).

### Transcription run-on and RNApol II-chromatin immunoprecipitation assays

The transcription run-on (TRO) assays of *PHO5-lacZ* fusions and *GAL1* were performed as in (Gómez-Herreros et al., 2012). *PHO5-lacZ* signals were normalized against the plasmid copy number as determined by Q-PCR, and by adapting the method described in (Skulj et al. 2008) to yeast cells. The total genomic DNA from plasmid-containing yeast was obtained and the Q-PCR signal of *lacZ* (amplicon corresponding to position 3 in Fig. 2A) was compared to the signal of the chromosomic *GAL1* gene. Q-PCR was carried out with SYBR Green Premix Ex Taq (Takara) in a Light Cycler 480 II (Roche).

The RNApol II ChIP experiments were run using anti-Rpb3 antibodies (ab81859; Abcam) as in (Begley et al. 2019). Sequences of the oligonucleotide employed in this work for the detection of *PHO5-lacZ* and *GAL1* by Q-PCR are the following: ATGCTCGTGACTTCTTGGCTC and AAAACGGCGAAACTGGTTTGG (position 1), GCACCGATCGCCCTTCCCAAC and CCAGGCAAAGCGCCATTCGCC (position 2), CGCGGCGACTTCCAGTTCAAC and AGATGGCGATGGCTGGTTTCC (position 3), CGGTCGTTGCAGAACATTATG and GATCTTCCTCACCGCAAACAG (*GAL1*).

## Abbreviations

SR: (Synthesis Rate)
NMD: (Non sense-Mediated Decay)
DR: (Degradation Rate)
[mRNA]_t_: (Total mRNA concentration)
PTC: (Premature Stop Codon)
HL: (mRNA Half Life)
RBP: (RNA-Binding Protein)
BF: (Buffering Factor)
RA: (mRNA amount or level)
GRO: (Genomic Run-On)
cDTA: (comparative Dynamic Transcriptome Analysis)
TRO: (Transcription Run-On)

## Acknowledgements

We thank A. Amon and M. Choder for their generous gift of yeast strains, and M. Choder and A. Singh for helpful discussion. This work was funded with grants from the Spanish Ministry of Economy and Competitiveness, the European Union (FEDER) [BFU2016-77728-C3-1-P to S. C.], [BFU2016-77728-C3-3-P to J.E.P-O] and [RED2018-102467-T to J.E.P-O and S.C.], the Regional Valencian Government [AICO2019/088 to J.E.P-O] and the Junta de Andalucía [BIO-271 to S.C.].

## Supplementary Information

**Table S1. List of the yeast strains used in this paper**.

**Table S2. Genomic data from the aneuploid mutant strains used in Figure 1A & C**.

**Table S3. Genomic data from the aneuploid strains used in Figure 1D**.

**Table S4. Genomic data from the *upf1* and BY4741 strains used in Figure 3**.

